# Allosteric Determinants of the SARS-CoV-2 Spike Protein Binding with Nanobodies: Examining Mechanisms of Mutational Escape and Sensitivity of the Omicron Variant

**DOI:** 10.1101/2021.12.22.473887

**Authors:** Gennady Verkhivker

## Abstract

Structural and biochemical studies have recently revealed a range of rationally engineered nanobodies with efficient neutralizing capacity against SARS-CoV-2 virus and resilience against mutational escape. In this study, we performed a comprehensive computational analysis of the SARS-CoV-2 spike trimer complexes with Nb6, VHH E and bi-paratopic VHH VE nanobodies. We combined atomistic dynamics and collective motions analysis with binding free energy scanning, perturbation-response scanning and network centrality analysis to examine mechanisms of nanobody-induced allosteric modulation and cooperativity in the SARS-CoV-2 spike trimer complexes with these nanobodies. By quantifying energetic and allosteric determinants of the SARS-CoV-2 spike protein binding with nanobodies, we also examined nanobody-induced modulation of escaping mutations and the effect of the Omicron variant on nanobody binding. The mutational scanning analysis supported the notion that E484A mutation can have a significant detrimental effect on nanobody binding and result in Omicron-induced escape from nanobody neutralization. Our findings showed that SARS-CoV-2 spike protein may exploit plasticity of specific allosteric hotspots to generate escape mutants that alter response to binding without compromising activity. The network analysis supported these findings showing that VHH VE nanobody binding can induce long-range couplings between the cryptic binding epitope and ACE2-binding site through a broader ensemble of communication paths that is less dependent on specific mediating centers and therefore may be less sensitive to mutational perturbations of functional residues. The results suggest that binding affinity and long-range communications of the SARS-CoV-2 complexes with nanobodies can be determined by structurally stable regulatory centers and conformationally adaptable hotspots that are allosterically coupled and collectively control resilience to mutational escape.

## 1. Introduction

SARS-CoV-2 infection is transmitted when the viral spike (S) glycoprotein binds to the host cell receptor ACE2, leading to the entry of S protein into host cells and membrane fusion [1,2]. The full-length SARS-CoV-2 S protein consists of amino (N)-terminal S1 subunit and carboxyl (C)-terminal S2 subunit where S1 is involved in the interactions with the host receptor and includes an N-terminal domain (NTD), the receptor-binding domain (RBD), and two structurally conserved subdomains (SD1 and SD2). Structural and biochemical studies established that the mechanism of virus infection may involve conformational transitions between distinct functional forms of the SARS-CoV-2 S protein in which the RBDs continuously switch between “down” and “up” positions [3–12]. The SARS-CoV-2 antibodies are divided into several main classes of which class 1 and class 2 antibodies target epitopes that overlap with the ACE2 binding site [13–15]. The body of structural and biochemical studies of the SARS-CoV-2 S complexes with different classes of potent antibodies targeting distinct binding epitopes of the S-RBD as well as various antibody cocktails and combinations have revealed multiple conformation-dependent epitopes, highlighting the link between conformational plasticity and adaptability of S proteins and capacity for eliciting specific binding and broad neutralization responses [16–36]. These studies have examined SARS-CoV-2 S binding with antibodies showing that combinations of antibodies can provide the efficient cross-neutralization effects through synergistic targeting of conserved and variable SARS-CoV-2 RBD epitope. Structural studies confirmed that the SARS-CoV-2 S protein can feature distinct antigenic sites and some specific antibodies may allosterically inhibit the ACE2 receptor binding without directly interfering with ACE2 recognition [30]. Optimally designed antibody cocktails simultaneously targeting different binding epitopes on the SARS-CoV-2 RBD demonstrated also improved resilience against mutational escape [37–42].

Nanobodies are much smaller than antibodies and thermostable, emerging as powerful therapies against SARS-CoV-2 [43]. An ultra-potent synthetic nanobody Nb6 neutralizes SARS-CoV-2 by stabilizing the fully inactive down S conformation preventing binding with ACE2 receptor [44]. A high-affinity trivalent nanobody mNb6-tri can simultaneously bind to all three RBDs and inhibit the interactions with the host receptor by occupying the binding site and locking the S protein in the inactive state [44]. The size-exclusion chromatography and mass spectrometry revealed high-affinity RBD-targeting nanobodies that efficiently neutralize SARS-CoV-2 by using several distinct and non-overlapping epitopes [45]. The revealed dominant epitope targeted by Nb20 and Nb21 nanobodies overlaps with the ACE2 binding site, showing that these nanobodies could competitively inhibit ACE2 binding and exploit structural mimicry to facilitate conformational changes that prematurely convert spike into a post-fusion state suppressing viral fusion [45]. Potent neutralizing nanobodies that resist circulating variants of SARS-CoV-2 by targeting novel epitopes were recently discovered [46]. The reported cryo-EM structures for different classes of nanobodies suggested mechanisms of high-affinity and broadly neutralizing activity by exploiting epitopes that are shared with antibodies as well as novel epitopes that are unique to the nanobodies [46]. The high affinity nanobodies against SARS-CoV-2 S protein refractory to common escape mutants and exhibiting synergistic neutralizing activity are characterized by proximal but non-overlapping epitopes showing that multimeric nanobody combinations can improve potency while minimizing susceptibility to escape mutations [47]. These studies identified a group of common resistant mutations in the dynamic RBM region (F490S, E484K, Q493K, F490L, F486S, F486L, and Y508H) that evade many individual nanobodies. Structural versatility of nanobody combinations that can effectively insulate the S-RBD accessible regions suggested a mechanism of resistance to mutational escape in which combining two nanobodies can markedly reduce the number of allowed substitutions to confer resistance and thereby elevate the genetic barrier for escape [47,48]. Using human VH-phage library and protein engineering several unique VH binders were discovered that recognized two separate epitopes within the ACE2 binding interface with nanomolar affinity [48]. Multivalent and bi-paratopic VH constructs showed markedly increased affinity and neutralization potency to the SARS-CoV-2 virus when compared to standalone VH domain [48]. Using saturation mutagenesis of the RBD exposed residues combined with fluorescence activated cell sorting for mutant screening, escape mutants were identified for five nanobodies and were mostly mapped to periphery of the ACE2 binding site with K417, D420, Y421, F486, and Q493 emerging as notable hotspots [49]. A wide range of rationally engineered nanobodies with efficient neutralizing capacity and resilience against mutational escape was recently unveiled that included the llama-derived nanobody VHH E bound to the ACE2-binding epitope and three alpaca-derived nanobodies VHHs U, V, and W that bind to a different cryptic RBD epitope [50]. Using X-ray crystallography and surface plasmon resonance-based binding competition this study showed that combinations of nanobodies targeting distinct epitopes can suppress the escape mutants resistant to individual nanobodies, while the biparatopic VHH EV and VE nanobodies with two antigen-binding sites appeared to be even more effective than pairs VHH E+U, E+V and E+W in preventing mutual escape [50–52]. Using single-domain antibody library and PCR-based maturation, two closely related and highly potent nanobodies, H11-D4 and H11-H4 were reported that recognize the same epitope immediately adjacent to and partly overlapping with the ACE2 binding region [53]. The crystal structures of these nanobodies bound to the S-RBD revealed binding to the same epitope, which partly overlaps with the ACE2 binding surface, explaining competitive inhibition of ACE2 interactions. These studies demonstrated that nanobodies may have potential clinical applications due to the increased neutralizing activity and robust protection against escape mutations of SARS CoV-2.

The high-affinity nanobody cocktails of two noncompeting nanobodies can neutralize both wild-type SARS-CoV-2 and the variants [54]. Neutralization of SARS-CoV-2 by low-picomolar and mutation-tolerant VHH nanobodies that bind synergistically to the opposite sides of the RBD produced a binding avidity effect unaffected by immune-escape mutants K417N/T, E484K, N501Y, and L452R [55]. The nanobody cocktails from camelid mice and llamas that neutralize SARS-CoV-2 variants showed a remarkable ability of multivalent nanobodies to combat escaping mutations through synchronized avidity between binding epitopes.^67^ In particular, picomolar nanobodiesNb12 and Nb30 revealed binding to a conserved RBD epitope outside of the ACE2-binding motif which is not accessible to human antibodies allowing for to combat escape mutations at E484 and N501 positions [56]. These studies suggested that nanobody mixtures and rationally engineered bi-paratopic nanobody constructs could offer a promising alternative to conventional monoclonal antibodies and may be advantageous for controlling a broad range of infectious variants while also suppressing the emergence of virus escape mutations. Furthermore, bi-paratopic nanobodies showed significant advantages compared to monoclonal antibodies, single nanobodies and nanobody cocktails by effectively leveraging binding avidity and allosteric cooperativity mechanisms in combating escape mutations.

Computer simulations and protein modeling played an important role in shaping up our understanding of the dynamics and function of SARS-CoV-2 glycoproteins [57–79]. All-atom molecular dynamics (MD) simulations of the full-length SARS-CoV-2 S glycoprotein embedded in the viral membrane, with a complete glycosylation profile were first reported by Amaro and colleagues, providing the unprecedented level of details and significant structural insights about functional S conformations [60]. A “bottom-up” coarse-grained (CG) model of the SARS-CoV-2 virion integrated data from cryo-EM, x-ray crystallography, and computational predictions to build molecular models of structural SARS-CoV-2 proteins assemble a complete virion model [61]. By providing valuable insights and establishing the blueprint for computational modeling, these studies paved the way for simulation-driven studies of SARS-CoV-2 spike proteins, also showing that conformational plasticity and the alterations of the SARS-CoV-2 spike glycosylation can synergistically modulate complex phenotypic responses to the host receptor and antibodies.

Multi-microsecond MD simulations of a 4.1 million atom system containing a patch of viral membrane with four full-length, fully glycosylated and palmitoylated S proteins allowed for a complete mapping of generic antibody binding signatures and characterization of the antibody and vaccine epitopes [62]. More recent extensive MD simulations and free energy landscape mapping studies of the SARS-CoV-2 S proteins and mutants detailed conformational changes and diversity of ensembles, further supporting the notion of enhanced functional and structural plasticity of S proteins [68–73]. Our recent studies combined coarse-grained and atomistic MD simulations with coevolutionary analysis and network modeling to present evidence that the SARS-CoV-2 spike protein function as allosterically regulated machine that exploits plasticity of allosteric hotspots to fine-tune response to antibody binding [74–79]. These studies showed that examining allosteric behavior of the SARS-CoV-2 pike proteins may be useful to uncover functional mechanisms and rationalize the growing body of diverse experimental data. Using MD simulations and protein stability analysis we recently examined binding of the SARS-CoV-2 RBD with single nanobodies Nb6 and Nb20, VHH E, a pair combination VHH E+U, a bi-paratopic nanobody VHH VE, and a combination of CC12.3 antibody and VHH V/W nanobodies [80]. This study provided a preliminary quantitative analysis of the escaping mutations on binding but was limited by consideration of only S-RBD regions

In the present work, we considerably expanded analysis of the SARS-CoV-2 S protein binding with nanobodies by performing a large number of high resolution coarse-grained (CG) simulations followed by full atomistic reconstruction for S protein trimer complexes with multivalent nanobodies Nb6, VHH E and bi-paratopic VHH VE nanobodies. We combined atomistic dynamics and collective motions analysis with binding free energy scanning, perturbation-response scanning and network modeling to examine mechanisms of nanobody-induced allosteric modulation and cooperativity in the SARS-CoV-2 S trimer complexes with nanobodies. By quantifying energetic and allosteric determinants of the SARS-CoV-2 S binding with nanobodies, we also examine nanobody-induced modulation of escaping mutations and the effect of the Omicron variant on nanobody binding. The results suggest that binding affinity and allosteric signatures of the SARS-CoV-2 complexes can be determined by a dynamic cross-talk between structurally stable regulatory centers and conformationally adaptable allosteric hotspots that collectively control resilience to mutational escape.

## 2. Results and Discussion

### 2.1. Conformational Dynamics and Collective Motions of the SARS-CoV-2 S Trimer Complexes: Nanobody-Induced Modulation of Flexibility and Escape Mutation Sites as Regulatory Hinges

We performed multiple CG simulations of the SARS-CoV-2 S trimer protein complexes with a panel of nanobodies (Figure 1) followed by all-atom reconstruction of trajectories to examine how structural plasticity of the RBD regions can be modulated by binding and determine specific dynamic signatures induced by different classes of nanobodies targeting distinct binding epitopes. All-atom MD simulations with the explicit inclusion of the glycosylation shield could provide a rigorous assessment of conformational landscape of the SARS-CoV-2 S proteins, such direct simulations remain to be technically challenging due to the size of a complete SARS-CoV-2 S system embedded onto the membrane. We combined CG simulations with atomistic reconstruction and additional optimization by adding the glycosylated microenvironment.

**Figure 1.**
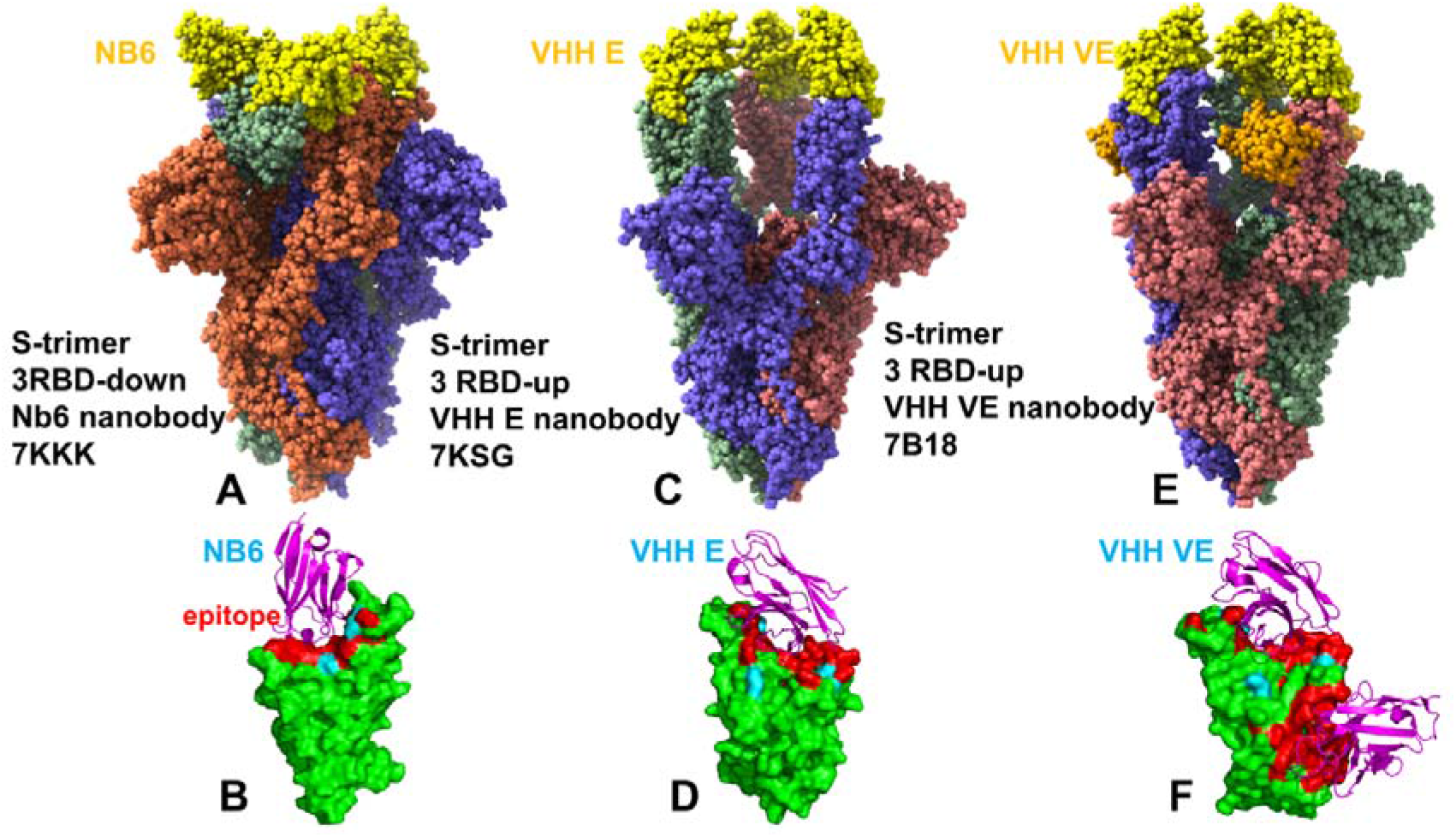
Cryo-EM structures of the SARS-CoV-2 S trimer complexes with a panel of nanobodies. (A) The structure of SARS-CoV-2 S trimer in the complex with Nb6 nanobody, pdb id 7KKK. Nb6 nanobodies are shown in yellow. (B) The S-RBD bound to Nb6. The S-RBD structures are shown in green surfaces. The binding epitope residues of the S-RBD bound structures are shown in red. (C) The SARS-CoV-2 S trimer in the complex with VHH E nanobody, pdb id 7KSG. VHH E is in yellow spheres. (D) The S-RBD bound to VHH E. (E) The SARS-CoV-2 S trimer in the complex with VHH VE nanobody, pdb id 7B18. VHH VE is in yellow/orange spheres. VHH E part in yellow and VHH V is in orange. The structures are shown in full spheres with protomers A,B,C colored in green, red and blue. The rendering of SARS-CoV-2 S structures was done using the interactive visualization program UCSF ChimeraX [81].

The results showed that Nb6 binding to the closed conformation of the S trimer can induce a more significant stabilization of the S-RBD and RBM residues (Figure 2A). A comparative analysis of the conformational flexibility profiles for the S trimer complexes with Nb6, VHH E, and VHH VE nanobody revealed stabilization of the interacting regions that was particularly strong in the complex with the VHH VE nanobody (Figure 2A). The RBD core α-helical segments (residues 349-353, 405-410, and 416-423) showed small thermal fluctuations in all complexes. The stability of the central ß strands (residues 354-363, 389-405, and 423-436) was especially pronounced in the S trimer complex with Nb6 nanobody (Figure 2A). A greater level of flexibility was seen for the S-RBD regions in the S trimer complexes with VHH E and VHH VE nanobodies (Figure 2A). It is evident that the conformational plasticity of the RBD-up conformations can be still maintained in the complexes with nanobodies. Although the VHH E epitope is very similar to that of other nanobodies such as Nb6, H11-D4, MR17, and SR4, VHH E binds in a specific orientation in which an extended β-hairpin conformation protrudes into the RBD binding site (Figure 2 B,C). Importantly, binding of the VHH VE nanobody to the cryptic epitope restricted mobility of the S2 subunit residues (Figure 2A). Conformational dynamics profiles reaffirmed stability of the α-helical segments in the RBD that are located near the cryptic binding epitope (residues 369-384) targeted by VHH V nanobody. These residues provide a stable anchoring platform for nanobodies binding to the cryptic epitope, allowing for modulation of RBD flexibility and optimization of binding interactions with the RBD binding epitope (Figure 2). We also noticed that VHH E and VHH VE can induce greater stability of the S2 region as compared to Nb6 binding, while these nanobodies also allow for more plasticity in the S1 regions.

**Figure 2.**
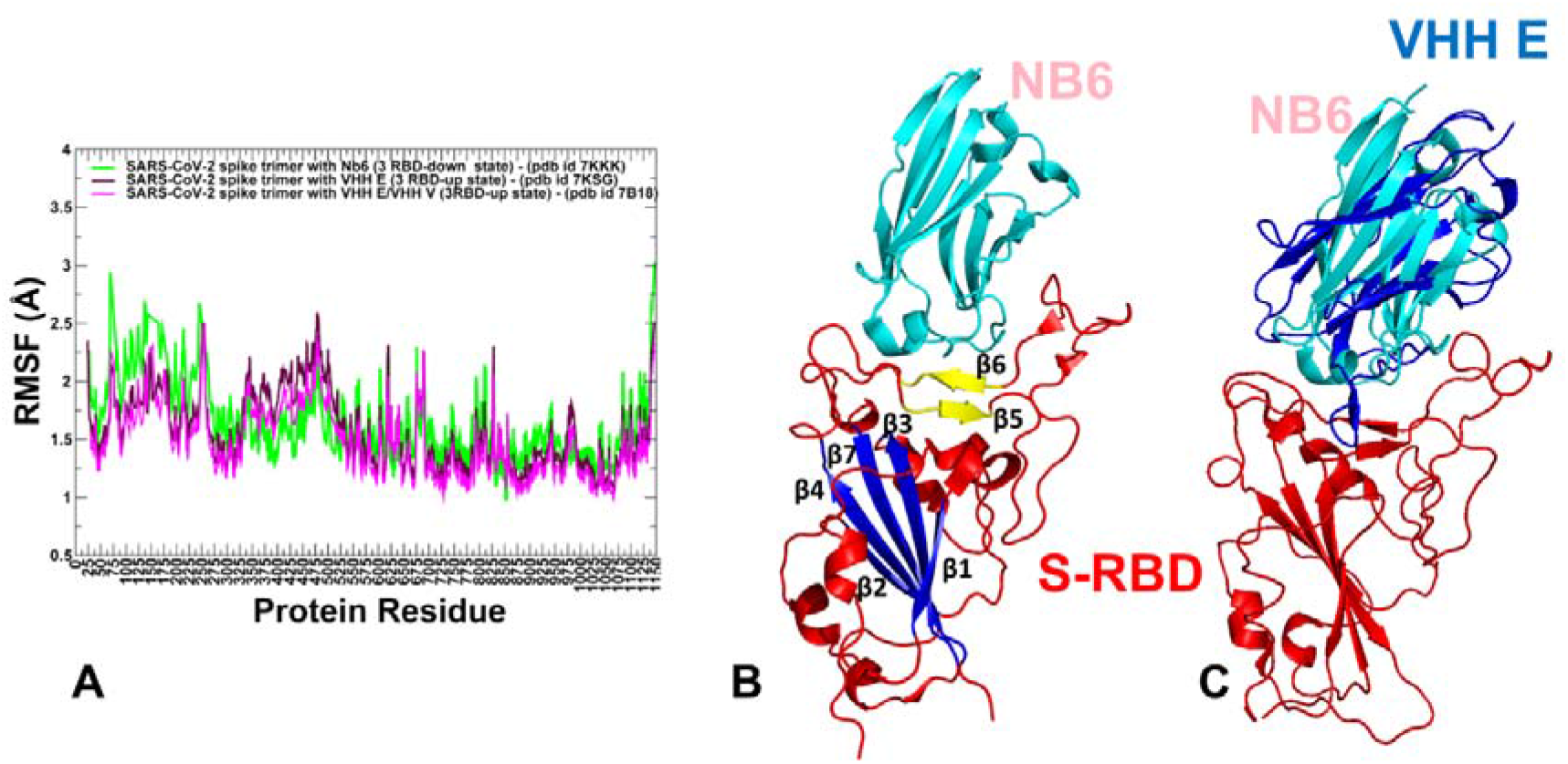
Conformational dynamics profiles of the SARS-CoV-2 S trimer complexes with nanobodies. (A) The root mean square fluctuations (RMSF) profiles obtained from simulations of the SARS-CoV-2 S trimer in the complex with Nb6 nanobody, pdb id 7KKK (in green lines), S trimer in the complex with VHH E nanobody, pdb id 7KSG (in maroon lines), and S trimer in the complex with VHH VE nanobody, pdb id 7B18 (in blue lines). (B) Structural organization of the S-RBD (shown in red ribbons). The central β strands (β1 to β4 and β7) (residues 354-358, 376-380, 394-403, 431-438, 507-516) are shown in blue. β5 and β6 strands (residues 451-454 and 492-495) are shown in yellow. The bound nanobody Nb6 is shown in cyan ribbons. (C) Superposition of Nb6 nanobody (in cyan ribbons) and VHH E nanobody (in blue ribbons). S-RBD is in red ribbons.

We found that the nanobody-induced stabilization of the 3 RBD-up conformation in the S trimer complexes with VHH E and VHH VE could still be balanced by appreciable level of intrinsic flexibility in the RBD-up conformations (Figure 2). Structural maps of the conformational dynamics profiles for the S-RBD complexes with Nb6 (Figure 3A), VHH E (Figure 3B) and VHH VE (Figure 3C) illustrated an appreciable mobility of the NTD and RBD residues in the 3-up complexes with VHH E and VHH E/VHH V nanobodies.

**Figure 3.**
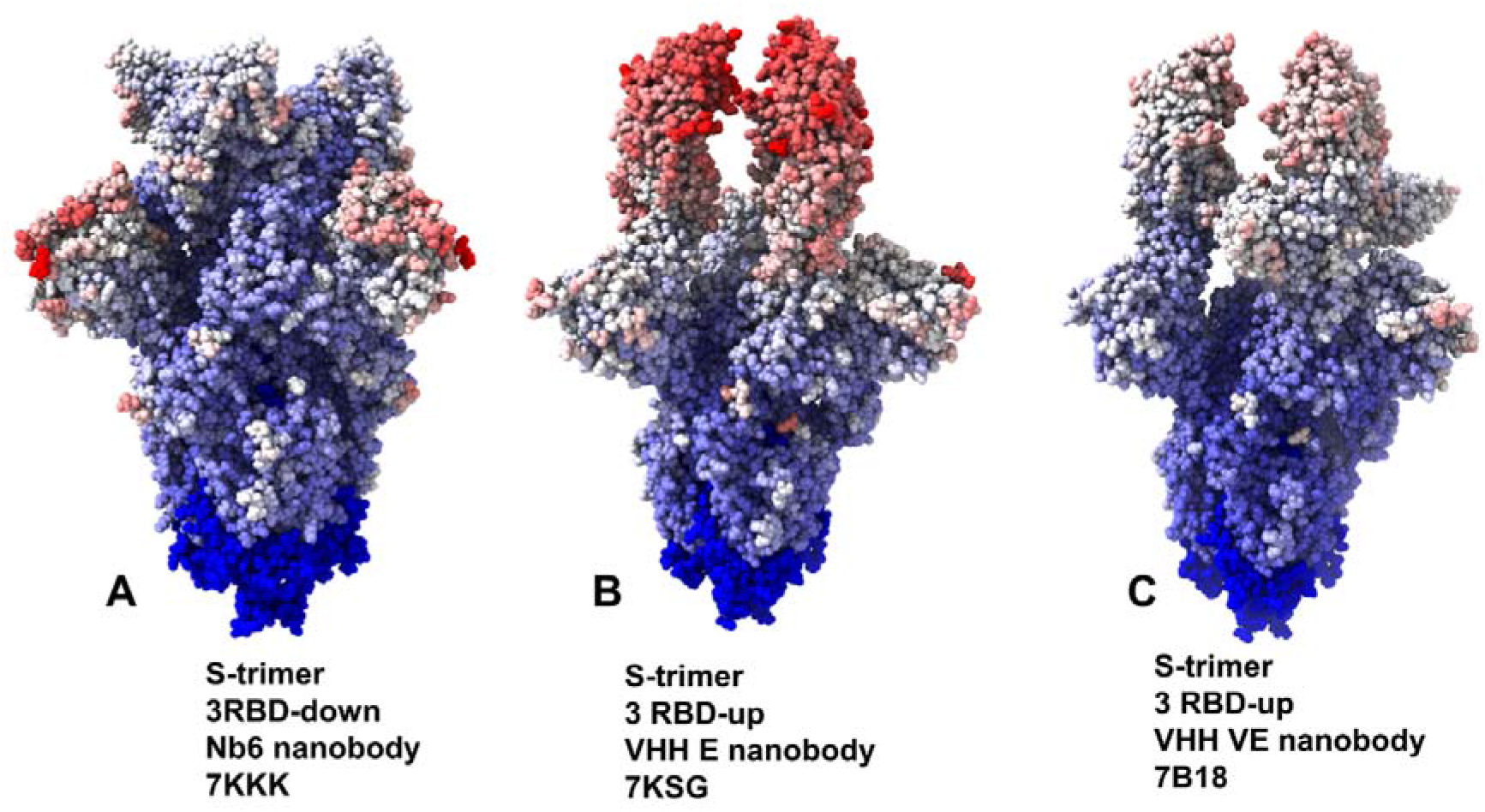
Structural maps of the conformational mobility profiles for the SARS-CoV-2 S protein variants. The dynamics maps for the SARS-CoV-2 S trimer in the complex with Nb6 nanobody, pdb id 7KKK (A), S trimer in the complex with VHH E nanobody, pdb id 7KSG (B), and S trimer in the complex with VHH VE nanobody, pdb id 7B18 (C). The structures are in sphere-based representation rendered using UCSF ChimeraX [81] with the rigidity-to-flexibility sliding scale colored from blue to red.

We characterized collective motions for the SARS-CoV-2 S-RBD complexes averaged over low frequency modes using principal component analysis (PCA) of the trajectories (Figure 4). The local minima along these profiles are typically aligned with the immobilized in global motions hinge centers, while the maxima correspond to the moving regions undergoing concerted movements leading to global changes in structure. The low-frequency ‘soft modes’ are often functionally important as mutations or binding can exploit and modulate protein movements along the pre-existing slow modes to induce allosteric transformations. The overall shape of the essential profiles in the SARS-CoV-2 S trimer complex with Nb6 showed suppressed movements of RBDs that are in the down position (Figure 4A,B). On the other hand, the profile displayed large functional displacements of the NTD regions. The immobilized hinge positions of the S trimer correspond to positions F318, L387, F429. The slow mode profile of the S trimer complex with Nb6 showed the reduced RBD mobility but the tip of the RBM loop (residues 473-483) remained mobile in functional dynamics. The sites of typical nanobody-escaping mutations (G447, Y449, L452, F490, Q493, Y508) correspond to the low mobility RBD regions in slow modes of the S trimer (Figure 4A,B).

**Figure 4.**
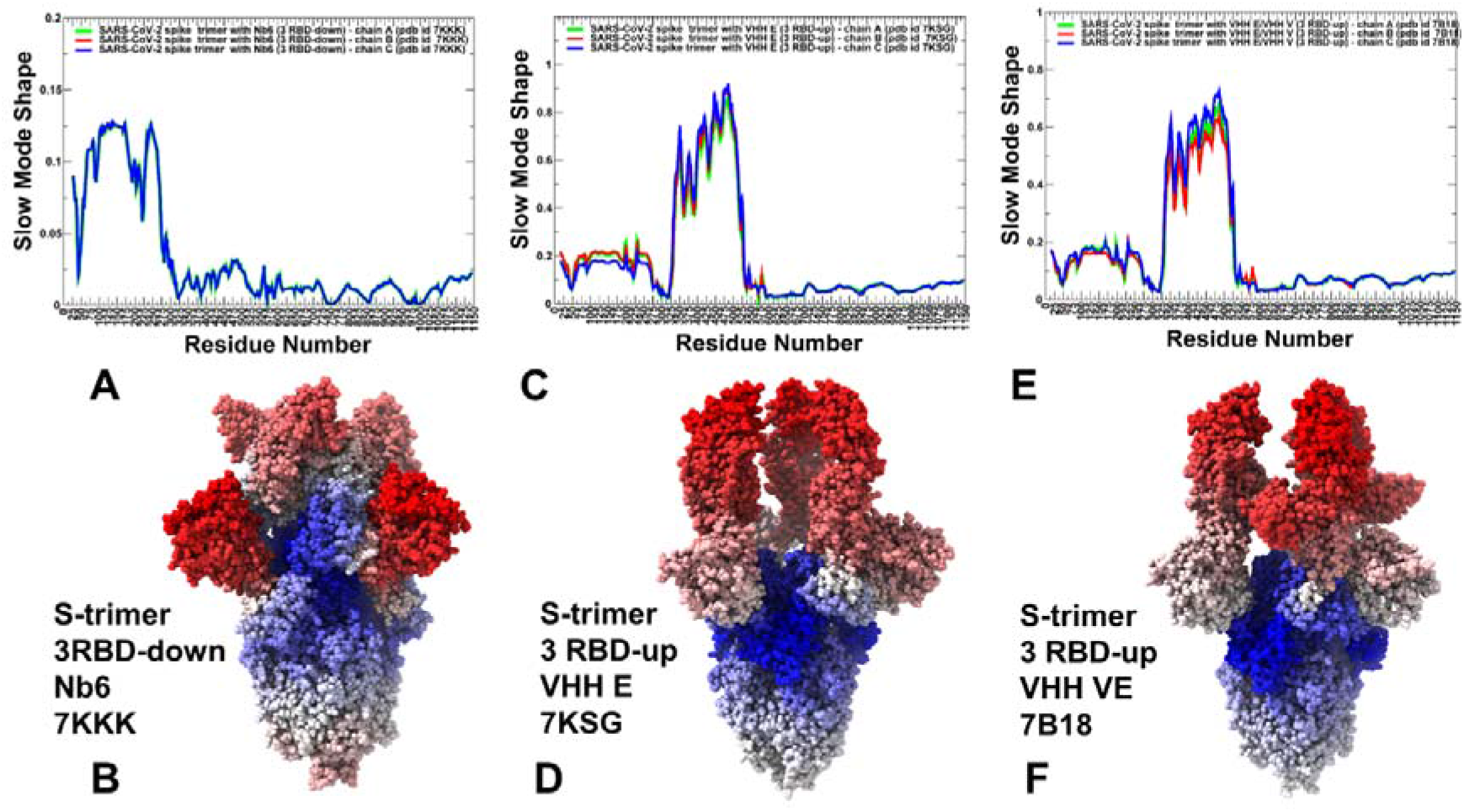
Functional dynamics of the SARS-CoV-2 S trimer structures. The essential mobility profiles are averaged over the first three major low frequency modes. The essential mobility profiles and structural maps for the SARS-CoV-2 S trimer in the complex with Nb6 nanobody, pdb id 7KKK (A,B), S trimer in the complex with VHH E nanobody, pdb id 7KSG (C,D), and S trimer in the complex with VHH VE nanobody, pdb id 7B18 (E,F). The structures on panels (B,D,F) are in sphere-based representation rendered using UCSF ChimeraX [81] with the rigidity-to-flexibility sliding scale colored from blue to red.

Although the RBD region harboring E484/F486 positions undergoes some functional motions in the slow modes, these movements are relatively moderate as compared to the NTD fluctuations that dominate collective dynamics. Nb6 binding could be severely compromised by the E484K mutation, while other sites of nanobody-escaping mutations are likely to be suppressed by the nanobody [44]. This may be partly explained based on the functional dynamics profiles in which most of these positions are immobilized by Nb6 binding, whereby the absence of functional motions could restrict the mutational escape potential. The fact that only the tip of the RBM region and E484/F486 remain more prone to changes could allow for E484K mutation to escape Nb6 binding and adopt a conformation evading efficient nanobody interactions.

The slow mode profile of the S trimer complex with VHH E nanobodies in which all RBDs are in the up position showed a clearly different pattern (Figure 4C,D). In this case, the RBDs correspond to moving regions. The rigid hinge centers are still located at conserved F318 and V534, F592 residues. Several local hinge positions are aligned with I358, A363, Y365, L387 in the RBD core due to constraints imposed by RBD interactions with NTD of the adjacent protomer. The local maxima of the slow mode profile in the RBD regions corresponded to V350, V369, S371, F377, K378, G447, Y449, L452, and 476-492 cluster (Figure 4C,D). Some of these functionally mobile residues are not involved in the interactions with VHH E nanobody (V350, V369, S371, F377, K378) and allow for conformational rearrangements of these flexible RBD regions. Instructively, some escaping mutations can target residues S375-Y380. In addition, nanobody binding can be partly escaped by mutations Y369H, S371P, F377L and K378Q/N even though these modifications are not currently circulating. Hence, the sites of escaping mutations are aligned with the functionally moving RBD regions which may experience functional displacements and affect the RBD conformation thereby reducing the efficiency of VHH E binding. The largest peaks in the slow mode profile is aligned with K417, F456 and RBM residues E484/F486 (Figure 4C,D). Movements of these positions may affect fidelity of nanobody binding and mutations in these positions, particularly E484K, can escape nanobody effect owing to the inherent functional plasticity in this region. This my contribute to a certain level of vulnerability shown by class 1 nanobodies (Nb6, VHH E) targeting the ACE2-binding site to mutations in K417 and E484 residues. Structural maps of the slow mode profiles for the S complex with VHH E (Figure 4D) illustrated the greater mobility of the RBM residues and plasticity of the binding epitope. A similar picture was observed for collective dynamics analysis of the S complex with VHH VE nanobody (Figure 4E,F). Our analysis indicated that VHH VE nanobody could modulate conformational dynamics without dramatically altering collective motions but rather fine-tune dynamic changes at the binding site. These findings are consistent with the experimental evidence showing that VHH E and VHH V nanobodies that target two independent epitopes can exploit S protein flexibility to activate the SARS-CoV-2 fusion machinery [50]. Although VHH VE binding can curtail flexibility of the S1 regions and impose structural constraints in the binding sites, functional RBD motions are still characteristic of the S complexes may contribute to mutational adaptation as sequences containing mutations in both interfaces were detected in the presence of VHHs E and V [50]. The results may explain why flexible RBD sites F486 and F490 are often featured as common sites of escape mutants that dominate the VHH E interface [50].

### 2.2. Mutational Scanning Identifies Structural Stability and Binding Affinity Hotspots in the SARS-CoV-2 Complexes and Explains Patterns of Nanobody-Escaping Mutations

By employ the conformational ensembles of the S trimer complexes with nanobodies we performed mutational scanning and computed binding free energy changes for studied SARS-CoV-2 S complexes with NB6, VHH E, and combination of VHH E/VHH V. In silico mutational scanning was done using BeAtMuSiC approach [82–84] This approach allows for accurate predictions of the effect of mutations on both the strength of the binding interactions and on the stability of the complex using statistical potentials and neural networks. This approach is showed a comparable performance and accuracy as physics-based FoldX potentials [85–88]. The adapted in our study BeAtMuSiC approach that was further enhanced through ensemble-based averaging of binding energy computations. The binding free energy ΔΔG changes were computed by averaging the results of computations over 1,000 samples obtained from simulation trajectories.

We first analyzed the mutational profiles for the S trimer 3-down complex with Nb6 (Figure 5). Mutational sensitivity analysis of the S binding with Nb6 showed results that were generally consistent with our earlier studies when using MD simulations of the S-RBD complex [80]. In the S trimer complex, however, a [single Nb6 molecule is positioned at the interface between two adjacent RBDs (Figure 1) [44]. The experimental studies suggested that a single Nb6 can stabilize two adjacent RBDs in the down state and prime the binding site for a second and third Nb6 molecule to stabilize the 3 RBD-down S conformation [44]. Mutational scanning of the S trimer revealed the binding energy hotspots in each protomer that are distributed through two interfaces, each interacting with a different Nb6 molecule (Figure 5). One of the interfaces corresponded to the cryptic binding RBD site where one Nb6 molecule interacts with N343, V367, S371, S373, V374, W436 hotspots (Figure 5). Our previous studies showed that highly conserved sites F374 and W436 are important coevolutionary centers that are often implicated in interactions with neutralizing antibodies [76,77]. The other Nb6 molecule binds to ACE2-binding site on the RBD where the key binding energy hotspots corresponded to hydrophobic residues Y449, L453, L455, F456, Y489, F490, G496 and Y505 (Figure 5). A number of these positions are also binding affinity hotspots for ACE2 as evident from deep mutagenesis scanning of SARS-CoV-2 interactions with the ACE2 host receptor [89–92]. The interaction pattern and similarity in the binding energy hotspots with ACE2 supported the notion of structural mimicry that may be efficiently exploited by Nb6 nanobody to competitively inhibit the ACE binding region.

**Figure 5.**
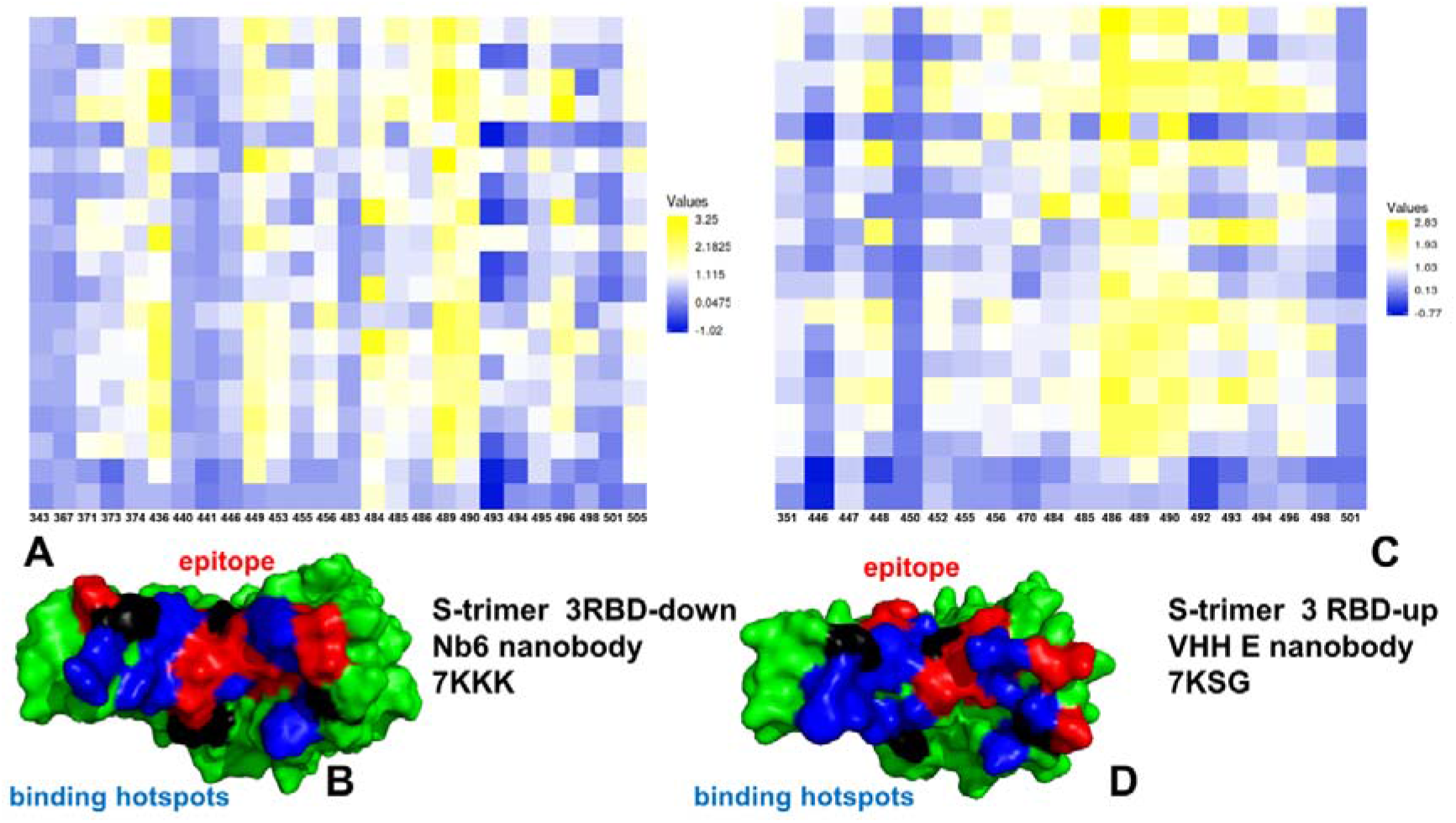
The mutational scanning heatmap for the SARS-CoV-2 S trimer complex with Nb6 nanobody, pdb id 7KKK (A,B) and VHH E nanobody, pdb id 7KSG (C,D). The binding energy hotspots correspond to residues with high mutational sensitivity. The heatmaps show the computed binding free energy changes for 19 single mutations on the binding epitope sites. The squares on the heatmap are colored using a 3-colored scale - from blue to yellow, with yellow indicating the largest destabilization effect. (B,D) Structural map of the binding epitopes and binding energy hotspots for Nb6 and VHH E. The S-RBD is shown in green surface. The epitope residues are shown in red and the binding energy hotspots from are shown in blue surface.

The mutational sensitivity map also shed some light on structure-functional role of sites targeted by common resistant mutations (F490S, E484K, Q493K, F490L, F486S, F486L, and Y508H) that evade many individual nanobodies [47]. Indeed, we found that E484, F486, and F490 positions can be sensitive to Nb6 binding (Figure 5). In particular, it was experimentally determined that Nb6 binding can be severely impeded by E484K mutation [50]. We specifically examined the effect of mutations present in the S-B.1.1.7 variant (N501Y) and S-B1.351 variant (K417N, E484K, N501Y on Nb6 and VHH E binding. It appeared that K417N and N501Y mutations only moderately affected nanobody binding. Somewhat more moderate but still noticeable destabilization changes can be induced in the S trimer complexes with VHH E nanobody upon mutations of L452 and E484 sites (Figure 5). Hence, these nanobody-escaping mutations center at highly antigenic sites. The moderate stability for sites of escaping mutations is consistent with the notion that virus tends to target positions where mutations would not appreciably perturb the RBD folding stability that is prerequisite for proper activity of spike protein and binding with the host receptor. By targeting dynamic and structurally adaptable hotspots such as E484, F486, and F490 that are relatively tolerant to mutational changes, virus tends to exploit conformational plasticity in these regions in eliciting specific escape patterns that would impair nanobody binding.

For the S trimer complex with VHH VE nanobody the binding footprint revealed several clusters of binding energy hotspots (Figure 6) targeting two different epitopes. The S-RBD hotspot residues correspond to Y449, L452, F456, F486, Y489, F490, and Y508 (Figure 5A). In agreement with the experiments [50], mutations at the VHH E interface Y449H/D/N, F490S, S494P/S, G496S, and Y508H produced destabilizing ΔΔG changes exceeding 2.0 kcal/mol (Figure 6). The binding epitope for VHH V is fairly large and includes Y369, N370, S371, A372, S373, F374, F377, L378, C379, Y380, G381, V382, S383 residues. The hotspot positions in the second cryptic epitope corresponded to the conserved and stable residues Y369, S371, F374, F377, C379, Y380 (Figure 6). The escaping mutations Y369H, S371P, F374I/V, T376I, F377L, and K378Q/N at the VHH U interface resulted in considerable destabilization losses (Figure 6). Hence, flexible RBD sites F486 and F490 are consistently featured as common binding energy hotspots for these complexes which may explain why escape mutants in these positions are known to dominate at the VHH E interface [50]. The results confirmed that nanobody combinations could alleviate the emergence and impact of escape mutants that target F456, F490 and Q493 residues.

**Figure 6.**
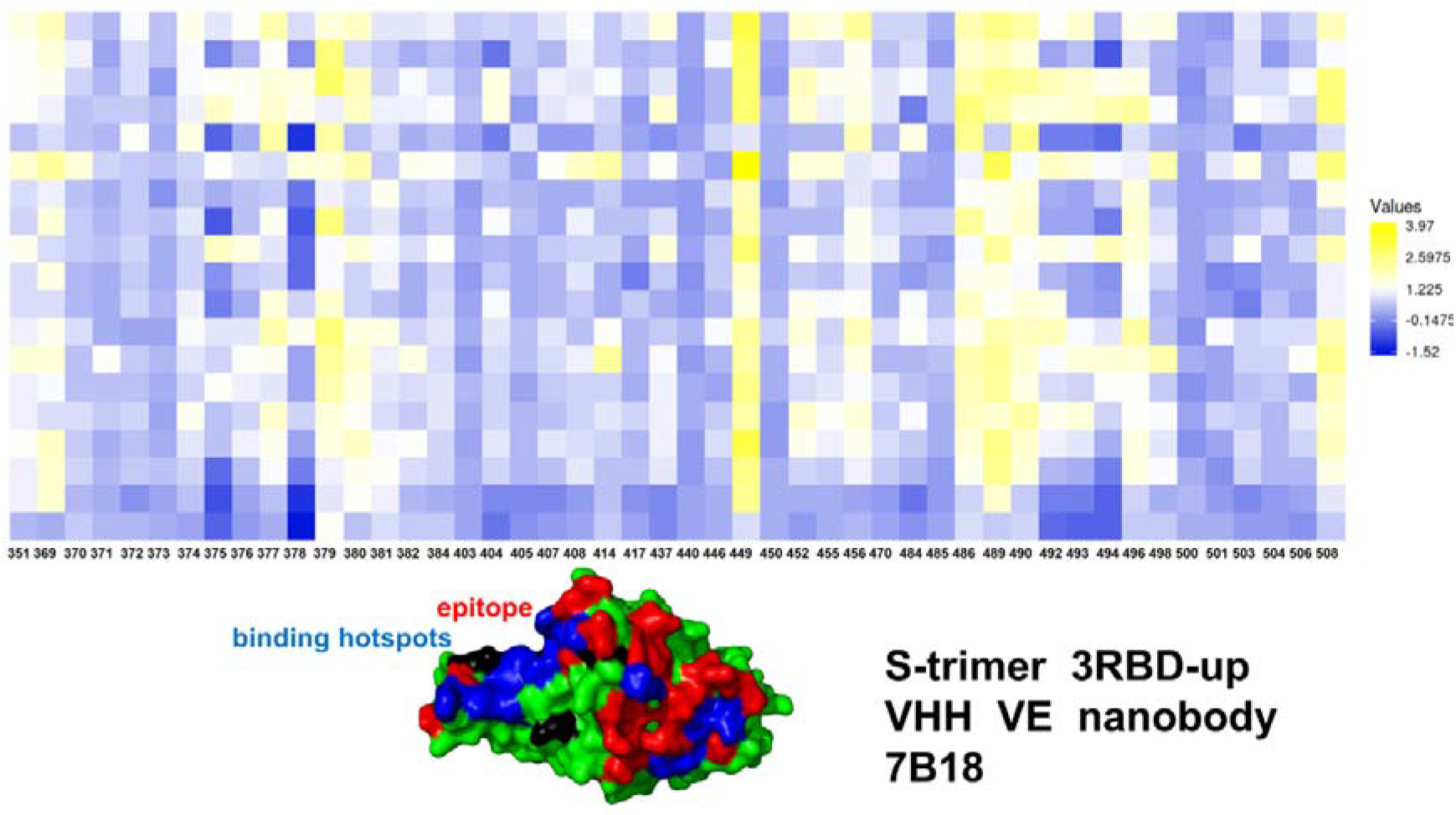
The mutational scanning heatmap for the SARS-CoV-2 S trimer complex with VHH VE nanobody, pdb id 7B18. The binding energy hotspots correspond to residues with high mutational sensitivity. The heatmaps show the computed binding free energy changes for 19 single mutations on the binding epitope sites. The squares on the heatmap are colored using a 3-colored scale - from blue to yellow, with yellow indicating the largest destabilization effect. Structural map of the binding epitopes and binding energy hotspots for VHH VE. The S-RBD is shown in green surface. The epitope residues are shown in red and the binding energy hotspots from are shown in blue surface.

We also examined the effect of Omicron mutations in the RBD (G339D, S371L, S373P, S375F, K417N, N440K, G446S, S477N, T478K, E484A, Q493R, G496S, Q498R, N501Y, Y505H) on binding of Nb6, VHH E, and VHH VE nanobodies (Figure 7). Importantly, some of the Omicron mutations could significantly affect Nb6 binding, particularly G446S, E484A, G496S, and Y50H modifications (Figure 7A). For VHH E binding, the largest binding affinity loss may result from E484A, Q493K, G496S, and N501Y mutations (Figure 7B). The important revelation of this analysis are noticeably smaller binding free energy changes in binding of VHH VE bi-paratopic antibody (Figure 7C). In this case, a noticeable binding affinity change may be induced by E484A, Q493K, and G496S mutations. In fact, these RBD mutations emerged as most sensitive hotspot cluster among Omicron RBD variant for nanobody binding (Figure 7). It was recently suggested that these mutations in the Omicron spike are compatible with usage of diverse ACE2 orthologues for entry and could amplify the ability of the Omicron variant to infect animal species [93]. Interestingly mutations in G446, L452, S477, T478, E484, F486, are associated with resistance to more than one monoclonal antibody and substitutions at E484 can confer broad resistance [94]. Several mutations at E484 position (E484A, E484G, E484D, and E484K) were discovered and each mutation has partial resistance to the convalescent plasma, indicating that E484 is also one of the dominant epitopes of spike protein [94,95]. The experimental studies also showed that E484 is the “Achilles’s heel” of several classes of nanobodies [45,46]. The mutational scanning analysis supported the notion that E484A mutation can have a significant detrimental effect on nanobody binding and result in Omicron-induced escape from nanobody neutralization.

**Figure 7.**
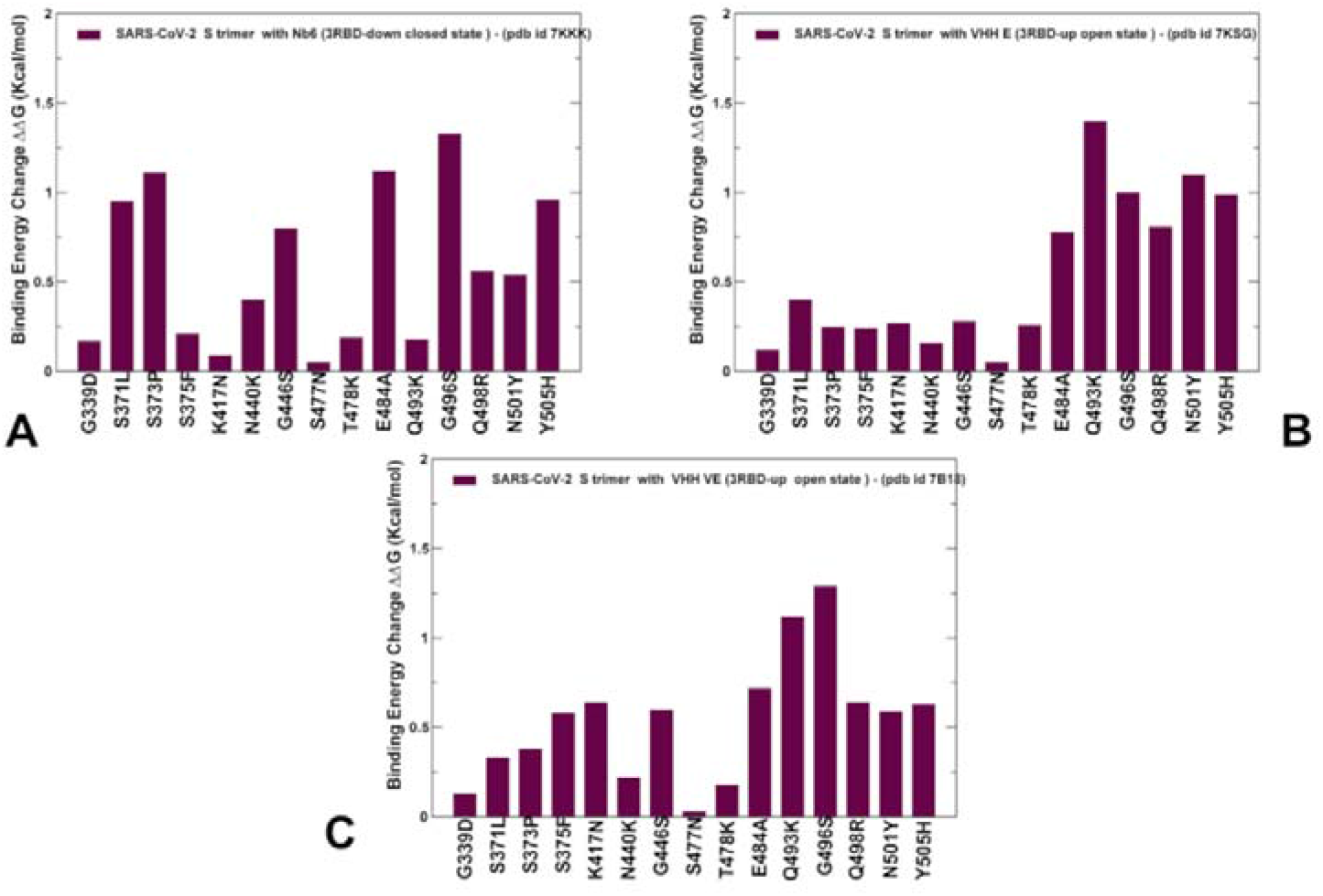
The mutational sensitivity analysis of the Omicron RBD mutations in the SARS-CoV-2 S trimer complexes with nanobodies. The binding free energy changes caused by Omicron RBD mutations on S trimer binding with Nb6 (A), VHH E (B) and VHH VE nanobody (C). The computed standard errors of the mean for the binding free energy changes based on different number of selected samples from a given trajectory (500, 1,000 samples) are 0.12-0.18 kcal/mol.

### 2.3 Perturbation Response Scanning of the SARS-CoV-2 S Complexes with Nanobodies Highlights Allosteric Role of Escaping Mutation Sites

Using the perturbation-response scanning (PRS) method [96–103] we quantified the allosteric effect of each residue in the SARS-CoV-2 complexes with a panel of studied nanobodies. The effector profiles estimate the propensities of a given residue to influence allosteric dynamic changes in other residues and are applied to identify regulatory hotspots of allosteric interactions as the local maxima along the profile. We propose that escaping variations could preferentially target structurally adaptable regulatory centers of collective movements and allosteric communications in the SARS-CoV-2 S complexes. To validate this hypothesis, we probed the allosteric effector potential of the S residues in complexes with studied nanobodies.

The PRS effector profile for the S-RBD residues in the complex with Nb6 showed a significant overlap with the complex with ACE2 (Figure 8A,B). In the complex with Nb6, several effector peaks corresponding to structurally stable RBD regions (residues 348-352, 400-406) as well as S371, S373, V374, W436 positions from the cryptic site involved in interactions with Nb6 nanobody. The largest effector values corresponded to RBD residues Q493, G496, L452, and Y508 (Figure 8A). Notably, a number of local maxima were also aligned with the sites of escaping mutations, particularly Y449, L452, L453, F490, L492, Q493 and Y508 positions (Figure 8A). Hence, these residues can exhibit a strong allosteric potential in the complex and function as effector hotspots of allosteric signal transmission (Figure 8A,B). In contrast, sites of circulating mutations K417, E484, and N501 belong to local minima of the profile which implies these residues as flexible sensors or transmitters of allosteric changes. This analysis also suggested that sites of escaping and circulating mutations may play a role in allosteric couplings of stable and flexible RBD regions that control signal propagation in the spike protein. While modifications of K417 and N501 residues appeared to trigger moderate changes in the binding affinity, the perturbations inflicted on these sites would have a significant effect on allosteric signaling in the complex. The results indicated that functional RBD sites may play complimentary roles in allosteric communications in the S complexes. While positions L452, Q493, G496 assume role of effector regulatory points that could dispatch allosteric signals though RBD regions, other functional sites such as more flexible E484, F486 and Y501 receive the signal and promote functional RBD movements. Structural mapping of allosteric effector hotspots for the S trimer complex with Nb6 nanobody revealed two clusters of residues: one cluster is in the S-RBD core region near the cryptic binding epitope and second cluster is near the RBM epitope (Figure 8B). These clusters form a network of functional centers that connects two binding epitopes and allow for signal transmission in the complex. It is particularly interesting given that Nb6 binds only to one of these binding epitopes. This suggests that allosteric effector centers in the RBD are allocated near the binding epitopes and are intrinsic to the S protein architecture. In this context, the pre-existing network of allosteric effector centers can be activated and modulated by nanobody binding that can exploit specific effector hotspots to allosterically propagate binding signal to other epitopes and functional regions. We also found that E484 site may be a critical effector hotspot for Nb6 binding. Allosteric versatility of this functional site could make it vulnerable to mutations which may alter collective dynamics and potentially be a driver of resistance to nanobodies. Indeed, mutations in the epitope centered on E484 position (F486, F490) were shown to strongly affect neutralization for different classes of nanobodies.

**Figure 8.**
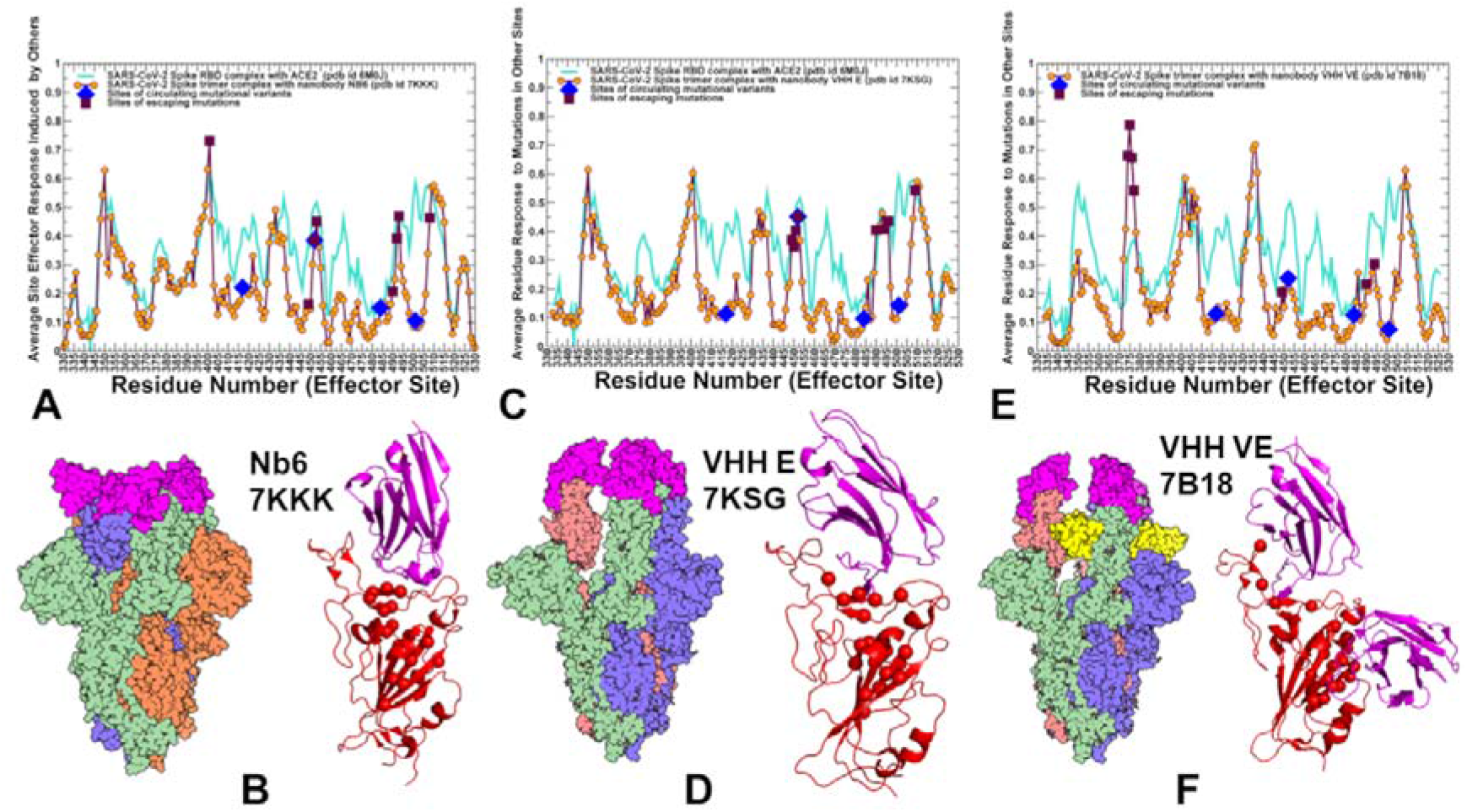
The PRS effector profiles for the SARS-CoV-2 S trimer complexes with Nb6 nanobody, pdb id 7KKK (A,B), VHH E nanobody, pdb id 7KSG (C,D) and VHH VE nanobody (E,F). The PRS effector profiles for the SARS-CoV-2 S complexes are shown in maroon-colored lines with orange-colored filled circles. For comparison, the PRS profiles are superimposed with the respective profiles for the S-RBD complex with ACE2 shown in cyan colored lines (pdb id 6M0J). The sites of escaping mutations for nanobody binding are indicated by maroon-colored filled squares and RBD sites K417, E484, and N501 targeted by global circulating variants are highlighted in blue-colored filled diamonds. (B,D,F) Structural maps of the allosteric effector hotspots corresponding to the local maxima of the PRS profiles. S trimer complexes are shown in sphere-based “flat” representation rendered using UCSF ChimeraX [81] with Nb nanobody (B) and VHH E nanobody shown in magenta. For VHH VE nanobody complex (F), VHH E is shown in magenta and VHH V is in yellow. Structural positions of allosteric effector centers are shown in red spheres and S-RBD is in red ribbons. The bound nanobodies are shown in magenta-colored ribbons.

The PRS profile of the S timer complex with VHH E nanobody (Figure 8C,D) featured RBD positions L452, Q493, G496, Q498, Y508 among pronounced peaks of the distribution, suggesting that these sites could function as regulatory sites of allosteric signaling in the complex. Similarly, to Nb6 complex, the structural map of the effector centers highlights a cluster near the cryptic binding site of the RBD core. The overall preservation of the topology and distribution of the allosteric effector centers is evident from our analysis, supporting the notion of pre-existing regulatory control points in the S protein.

Instructively, the PRS profile for the S complex with VHH VE nanobody that binds to two different binding sites revealed a partial redistribution of the allosteric centers (Figure 8E,F). In this case, the dominant sharp peak corresponded to a cluster of residues (S371, S373, V374, F377, K378) from the cryptic site that interact with VHH V. Smaller local peaks are associated with the RBD positions from the ACE2-binding site, primarily Q493, Q498, andY508 (Figure 8E). As a result, VHH VE binding could shifts the distribution towards allosteric sites from the cryptic binding site that regulate signal propagation in the S complex, while functional residues from the RBM binding site may serve as sensors of the binding signal. The diminished dependency of allosteric signaling induced by VHH VE nanobody on the common sites of escaping mutations may be related to the effects of multimeric nanobody combinations that allow for reduction of susceptibility to escape mutations. This suggests a plausible mechanism by which bi-paratopic nanobodies can leverage dynamic couplings to synergistically inhibit distinct binding epitopes and suppress mutational escape. To summarize, perturbation-based scanning results revealed allosteric role of functional sites targeted by escaping mutations and Omicron variant. Collectively, our findings suggested that SARS-CoV-2 S protein may exploit plasticity of specific allosteric hotspots to generate escape mutants that alter response to binding without compromising activity.

### 2.4 Network Centrality Analysis of Global Mediating Centers in the SARS-CoV-2 Complexes with Nanobodies Identifies Clusters of Allosteric Hotspots Targeted by Escaping Mutations

Network-centric models of protein structure and dynamics can allow for a more quantitative analysis of allosteric changes, identification of regulatory control centers and mapping of allosteric communication pathways. The residue interaction networks in the SARS-CoV-2 spike trimer structures were built using a graph-based representation of protein structures [104,105] in which residue nodes are interconnected through dynamic correlations [106]. By employing network centrality calculations for the equilibrium ensembles of the SARS-CoV-2 S trimer complexes with nanobodies [107,108], we computed ensemble-averaged distributions of the short path residue centrality (Figure 9). This network metric was used to identify mediating centers of allosteric interactions in the SARS-CoV-2 complexes. In the context of the network-based centrality analysis residues mediating a significant number of shortest pathways between all possible residue pairs in the system are identified by higher betweenness centrality. Residues with high betweenness centrality typically act mediating centers for the ensemble of optimal allosteric communication pathways.

**Figure 9.**
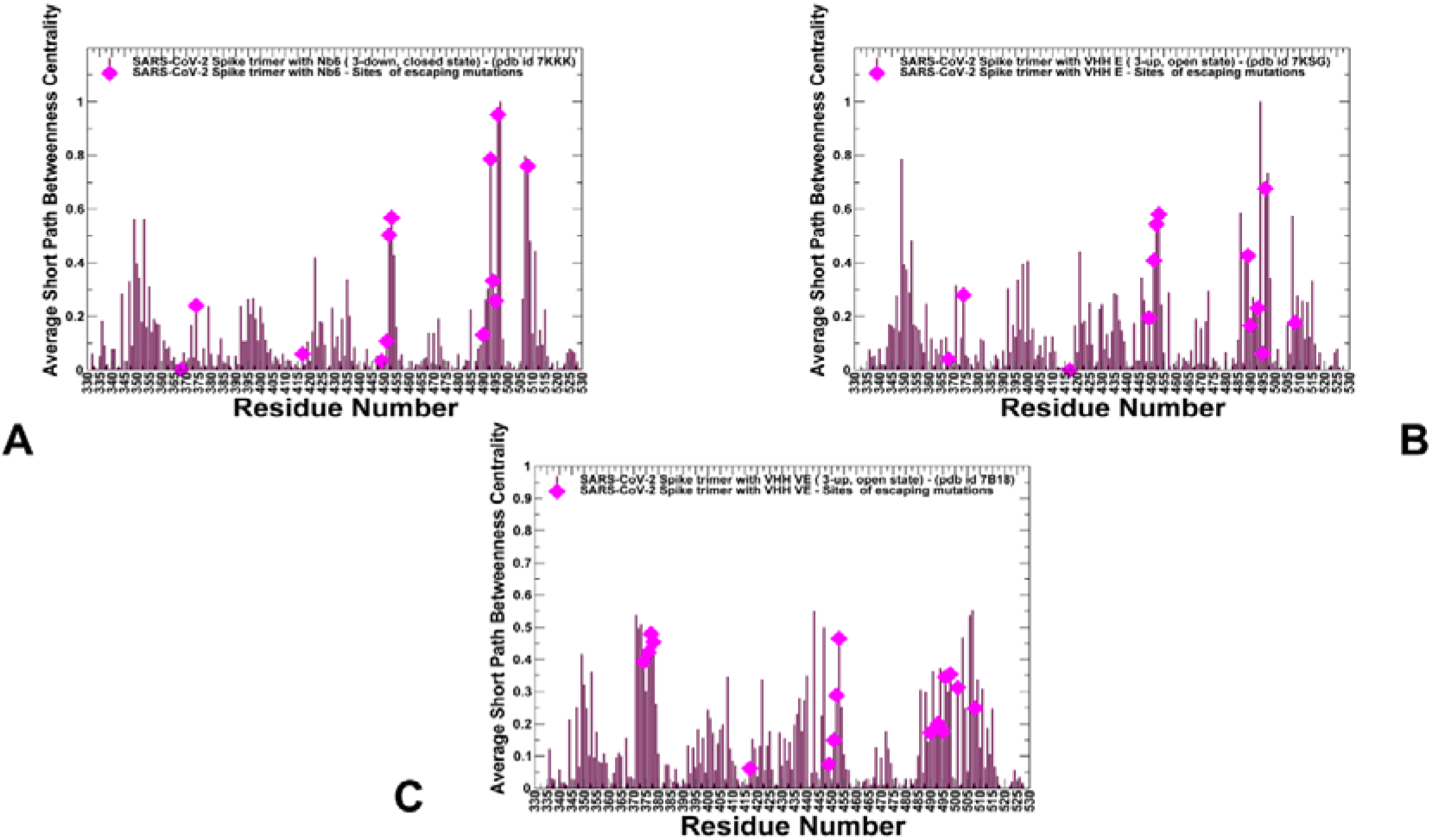
Network centrality analysis of the SARS-CoV-2 S trimer complexes with nanobodies. (A) The ensemble-averaged short path betweenness centrality for the SARS-CoV-2 S trimer in the complex with Nb6 nanobody, pdb id 7KKK (A), S trimer in the complex with VHH E nanobody, pdb id 7KSG (B), and S trimer in the complex with VHH VE nanobody, pdb id 7B18 (C). The residue-based profiles are shown for the S trimers are shown in maroon-colored filled bars. The sites of escaping mutations for nanobody binding are highlighted in blue-colored filled diamonds.

The network centrality profiles revealed several characteristic cluster peaks that are shared among complexes (Figure 9). However, nanobody binding can modulate this distribution and change the relative contribution of mediating centers. In the S trimer complexes with N6 and VHH E nanobodies that target the ACE2-binding sites, we observed the largest peak localized in the cluster of F490, L492, Q493, G496, Q498 and Y508 positions residues (Figure 9A,B). The second peak is aligned with Y449, L452, L453, and L455 RBD positions. In network terms, this implies that allosteric signaling in the S complexes with Nb6 and VHH E can be mediated by these sites that serve as central communication hubs. As a result, mutations in these positions and loss of interactions can affect not only the local structural environment of the mutated sites but also impact global network organization of the system. Strikingly, a significant number of these mediating centers corresponded to residues involved in the Omicron variant. Hence, multiple Omicron RBD mutations (such as Q493R, G496S, Q498R, N501Y, Y505H) may have a measurable effect on allosteric couplings in the complexes with Nb6 and VHH E nanobodies, which would likely render some level of resistance to nanobody-induced neutralization.

In some contrast, in the S complex with bi-paratopic VHH VE nanobody, a partial redistribution of the network centrality distribution was detected, pointing to the reduced peaks in the RBD residues from the ACE2-binding site, while showing a moderate centrality for S-RBD core residues from the cryptic site (S371, F374, S375, F377, C379, Y380). The observed modulation of high centrality peaks and broadening of the distribution showed that many residues feature a moderate level of centrality. As result, VHH VE nanobody binding can induce long-range couplings between the cryptic binding epitope and ACE2-binding site through a broader ensemble of communication paths that is less dependent on specific mediating centers and therefore may be less sensitive to mutational perturbations of functional residues. This suggests a plausible mechanism by which bi-paratopic nanobodies can leverage dynamic couplings to synergistically inhibit distinct binding epitopes and suppress mutational escape.

## 3. Materials and Methods

### 3.1 Structure Preparation and Analysis

All structures were obtained from the Protein Data Bank [109,110]. During structure preparation stage, protein residues in the crystal structures were inspected for missing residues and protons. Hydrogen atoms and missing residues were initially added and assigned according to the WHATIF program web interface [111,112]. The structures were further pre-processed through the Protein Preparation Wizard (Schrödinger, LLC, New York, NY) and included the check of bond order, assignment and adjustment of ionization states, formation of disulphide bonds, removal of crystallographic water molecules and co-factors, capping of the termini, assignment of partial charges, and addition of possible missing atoms and side chains that were not assigned in the initial processing with the WHATIF program. The missing loops in the studied cryo-EM structures of the SARS-CoV-2 S protein were reconstructed and optimized using template-based loop prediction approaches ModLoop [113], ArchPRED server [114] and further confirmed by FALC (Fragment Assembly and Loop Closure) program [115]. The side chain rotamers were refined and optimized by SCWRL4 tool [116]. The conformational ensembles were also subjected to all-atom reconstruction using PULCHRA method [117] and CG2AA tool [118] to produce atomistic models of simulation trajectories. The protein structures were then optimized using atomic-level energy minimization with a composite physics and knowledge-based force fields as implemented in the 3Drefine method [119]. The atomistic structures from simulation trajectories were further elaborated by adding N-acetyl glycosamine (NAG) glycan residues and optimized.

### 3.2 Coarse-Grained Simulations

Coarse-grained (CG) models are computationally effective approaches for simulations of large systems over long timescales. We employed CABS-flex approach that efficiently combines a high-resolution coarse-grained model and efficient search protocol capable of accurately reproducing all-atom MD simulation trajectories and dynamic profiles of large biomolecules on a long time scale [120–126]. In this high-resolution model, the amino acid residues are represented by Cα, Cβ, the center of mass of side chains and another pseudoatom placed in the center of the Cα-Cβ pseudo-bond. In this model, the amino add residues are represented by Cα, Cβ, the center of mass of side chains and the center of the Cα-Cα pseudo-bond. The CABS-flex approach implemented as a Python 2.7 object-oriented standalone package was used in this study to allow for robust conformational sampling proven to accurately recapitulate all-atom MD simulation trajectories of proteins on a long time scale [124–126]. Conformational sampling in the CABS-flex approach is conducted with the aid of Monte Carlo replica-exchange dynamics and involves local moves of individual amino acids in the protein structure and global moves of small fragments [136]. A total of 1,000 independent CG-CABS simulations were performed for each of the studied systems. In each simulation, the total number of cycles was set to 10,000 and the number of cycles between trajectory frames was 100. MODELLER-based reconstruction of simulation trajectories to all-atom representation provided by the CABS-flex package was employed to produce atomistic models of the equilibrium ensembles for studied systems [124]. We also performed principal component analysis (PCA) of reconstructed trajectories derived from CABS-CG simulations using the CARMA package [127].

### 3.3 Mutational Scanning and Sensitivity Analysis

We conducted mutational scanning analysis of the binding epitope residues for the SARS-CoV-2 S protein complexes. Each binding epitope residue was systematically mutated using all 19 substitutions and corresponding protein stability changes were computed. BeAtMuSiC approach [82–84] was employed that is based on statistical potentials describing the pairwise inter-residue distances, backbone torsion angles and solvent accessibilities, and considers the effect of the mutation on the strength of the interactions at the interface and on the overall stability of the complex. The binding free energy of protein-protein complex can be expressed as the difference in the folding free energy of the complex and folding free energies of the two protein binding partners:

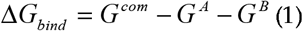

The change of the binding energy due to a mutation was calculated then as the following

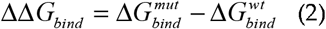

We leveraged rapid calculations based on statistical potentials to compute the ensemble-averaged binding free energy changes using equilibrium samples from simulation trajectories. The binding free energy changes were computed by averaging the results over 1,000 equilibrium samples for each of the studied systems.

### 3.4 Perturbation Response Scanning

Perturbation Response Scanning (PRS) approach [96–103] follows the protocol originally proposed by Bahar and colleague [98,99] and was described in detail in our previous studies [75,76]. In brief, through monitoring the response to forces on the protein residues, the PRS approach can quantify allosteric couplings and determine the protein response in functional movements. In this approach, it 3N × 3*N* Hessian matrix ***H*** whose elements represent second derivatives of the potential at the local minimum connect the perturbation forces to the residue displacements. The 3N-dimensional vector **Δ*R*** of node displacements in response to 3N-dimensional perturbation force follows Hooke’s law ***F*** = ***H * ΔR***. A perturbation force is applied to one residue at a time, and the response of the protein system is measured by the displacement vector Δ***R***(*i*) = ***H*^-1^*F*^(*i*)^** that is then translated into *N×N* PRS matrix. The second derivatives matrix ***H*** is obtained from simulation trajectories for each protein structure, with residues represented by *C_α_* atoms and the deviation of each residue from an average structure was calculated by Δ***R**_j_*(*t*) = ***R**_j_*(*t*) – 〈***R**_j_*(*t*)〉, and corresponding covariance matrix C was then calculated by Δ**R**Δ**R**^*T*^. We sequentially perturbed each residue in the SARS-CoV-2 spike structures by applying a total of 250 random forces to each residue to mimic a sphere of randomly selected directions. The displacement changes, Δ***R^i^*** is a 3*N*-dimensional vector describing the linear response of the protein and deformation of all the residues.

Using the residue displacements upon multiple external force perturbations, we compute the magnitude of the response of residue *k* as 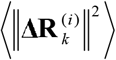 averaged over multiple perturbation forces **F**^(*i*)^, yielding the *ik*^th^ element of the *N×N* PRS matrix. The average effect of the perturbed effector site *i* on all other residues is computed by averaging over all sensors (receivers) residues *j* and can be expressed as〈(Δ***R^i^***)^2^)〉_*effector*_. The effector profile determines the global influence of a given residue node on the perturbations in other protein residues and can be used as proxy for detecting allosteric regulatory hotspots in the interaction networks. In turn, the *j*^th^ column of the PRS matrix describes the sensitivity profile of sensor residue *j* in response to perturbations of all residues and its average is denoted as 〈(Δ***R^i^***)^2^)〉_*sensor*_. The sensor profile measures the ability of residue *j* to serve as a receiver of dynamic changes in the system.

## 4. Conclusions

In this study, we performed a comprehensive computational analysis of the SARS-CoV-2 S trimer complexes with Nb6, VHH E and bi-paratopic VHH VE nanobodies. We combined atomistic dynamics and collective motions analysis with binding free energy scanning, perturbation-response scanning and network centrality analysis to examine mechanisms of nanobody-induced allosteric modulation and cooperativity in the SARS-CoV-2 S trimer complexes with nanobodies. By quantifying energetic and allosteric determinants of the SARS-CoV-2 S binding with nanobodies, we also examined nanobody-induced modulation of escaping mutations and the effect of the Omicron variant on nanobody binding. The mutational scanning analysis supported the notion that E484A mutation can have a significant detrimental effect on nanobody binding and result in Omicron-induced escape from nanobody neutralization. The results suggested that by targeting structurally adaptable hotspots such as E484, F486, and F490 that are relatively tolerant to mutational changes, virus tends to exploit conformational plasticity in these regions in eliciting specific escape from nanobody binding. Using PRS analysis, we found that escaping mutational variant could preferentially target structurally adaptable regulatory centers of collective movements and allosteric communications in the SARS-CoV-2 S complexes. We suggested that reduced dependency of allosteric signaling induced by VHH VE nanobody on the common sites of escaping mutations may be related to the effects of multimeric nanobody combinations that allow for reduction of susceptibility to escape mutations. Our findings showed that SARS-CoV-2 S protein may exploit plasticity of specific allosteric hotspots to generate escape mutants that alter response to binding without compromising activity. The network analysis supported these findings showing that VHH VE nanobody binding can induce long-range couplings between the cryptic binding epitope and ACE2-binding site through a broader ensemble of communication paths that is less dependent on specific mediating centers and therefore may be less sensitive to mutational perturbations of functional residues. The results suggest that binding affinity and long-range communications of the SARS-CoV-2 complexes with nanobodies can be determined by structurally stable regulatory centers and conformationally adaptable hotspots that are allosterically coupled and collectively control resilience to mutational escape.

## Author Contributions

Conceptualization, G.V.; methodology, G.V.; software, G.V.; validation, G.V.; formal analysis, G.V.; investigation, G.V.; resources, G.V.; data curation, G.V.; writing—original draft preparation, G.V.; writing—review and editing, G.V.; visualization, G.V.; supervision, G.V.; project administration, G.V.; funding acquisition, G.V. All authors have read and agreed to the published version of the manuscript.

## Acknowledgments

The author thanks Schmid College of Science and Technology, Chapman University for providing computing resources at the Keck Center for Science and Engineering

